# Error-driven changes in hippocampal representations accompany flexible re-learning

**DOI:** 10.1101/2025.05.20.655046

**Authors:** P. Dylan Rich, Stephan Y. Thiberge, Nathaniel D. Daw, David W. Tank

## Abstract

Flexible behavior requires both the learning of new associations, and the suppression of previous ones, but how neural circuits achieve this balance remains unclear. Here we show that continuous changes in hippocampal representations, known as drift, may facilitate this process. We used voluntary head-fixation and calcium imaging to record from CA1 in rats during an odor-guided navigation task that required frequent re-learning. We found systematic representational changes over the course of the multi-hour sessions that were increased following errors. A simple neural network model revealed that such error-driven drift can enable flexible re-learning by allowing new associations to form from new neural patterns. A consequence of this is that previous associations are maintained in latent synaptic weights. These findings reconcile the apparent tension between representational drift and stable memory storage, demonstrating how dynamic neural codes could support both flexible behavior and lasting memories.

## Introduction

Humans and animals must learn what to do in new situations. Such learning is thought to rely on the formation and strengthening of associations between neural representations of current states or situations and corresponding downstream actions that lead to a desired outcome (Brown and Sharp, 1995; McNaughton and Morris, 1987). The hippocampus is believed to play a central role in this process by providing a neural representation of states or situations; cells in the hippocampus code for locations (O’Keefe and Dostrovsky, 1971), collectively discriminate between different contexts (Leutgeb et al., 2004; McKenzie et al., 2014; Muller and Kubie, 1987), task states (Aronov and Tank, 2014; Pastalkova et al., 2008; Sun et al., 2020) and abstract quantities (Nieh et al., 2021). Moreover, the reactivation of hippocampal cells is sufficient to elicit the recall of previously learned behaviors (Liu et al., 2012; Robinson et al., 2020).

Although initially considered to be stable (Thompson and Best, 1990), spatial representations in the hippocampus change over time, even with the same conditions are revisited (Kentros et al., 2004; Manns et al., 2007; Ziv et al., 2013). This phenomenon, termed representational drift, provides a challenge for a state representation view of the hippocampus’ function, which tends to presuppose a static neural representation that can allow the retrieval of learned information. How the dynamic nature of hippocampal representation relates to its role in learning and memory remains to be explored.

One view is that representational drift is undesirable, a by-product of synaptic turn-over (Mongillo et al., 2017), and a problem for stable behavior (Driscoll et al., 2022), with mechanisms proposed for how downstream circuits can compensate for drift (Rule and O’Leary, 2022). An alternative view is that drift is useful, and that changes in representations are required for ongoing learning (Driscoll et al., 2022; Micou and O’Leary, 2023) and dynamic memory encoding (Mau et al., 2020). Computational models of memory function such as the temporal context model (Howard and Kahana, 2002) posit that an ongoing, changing neural representation links events that occur close in time (Cai et al., 2016; Rubin et al., 2015). A dynamic representation has also been proposed to support the adaptive learning rates that can be useful in volatile environments (Razmi and Nassar, 2022).

Although the idea that representational changes exist to accommodate changes in an environment and enable learning is intuitive, evaluating this hypothesis rigorously has been hampered by limitations in existing studies where drift has been observed. In order to effectively map place cells, animals are typically engaged in simple tasks, such as foraging, that do not have an explicit learning component that is under experimental control. Behavioral factors, such as running speed, that are known to modulate hippocampal responses (McNaughton et al., 1983) are often uncontrolled and may explain aspects of drift (Liberti et al., 2022; Sadeh and Clopath, 2022). Finally, drift is often assessed across days or sessions (Geva et al., 2023; Khatib et al., 2023), masking how the short timescale elements of the change (Bittner et al., 2017; Priestley et al., 2022) may relate to specific learning events.

To shed light on these issues, we recorded hippocampal activity while animals were engaged in a task where learning and re-learning could be examined multiple times within a session. We used voluntary head-fixation (Rich et al., 2024) and two-photon calcium imaging to record the activity of neurons in hippocampal region CA1 during the stimulus presentation epoch of an odor-cued navigation task. During this period animals’ heads were in the same position, allowing unambiguous attribution of neural firing differences to cognitive variables (Igarashi et al., 2014; MacDonald et al., 2013). Crucially, the spatial configuration of the maze animals navigated was changed multiple times per session, requiring animals to periodically adapt to new contingencies.

We found the task representation in CA1 was driven by the cue stimulus, changed throughout the session, and that these changes were increased following trials when the animal made a navigational error. To understand the consequences of drift in CA1 to a downstream circuit, we simulated a simple feed-forward network that had to perform the same task and adapt to changing odor-action associations. The model illustrated the advantage of the evolving task state representation for both learning new associations and suppression of previous, irrelevant ones (Razmi and Nassar, 2022). However, previous rules were not entirely lost, but, as a consequence of the ongoing drift, were preserved in latent synaptic weights. For this reason, such a model suggests that representational drift could serve as a common mechanism bridging two processes that have typically been thought to have separate neural and computational substrates (Yu et al., 2021): both incremental adaptive learning (Razmi and Nassar, 2022) and latent state retrieval.

## Results

We trained animals to perform a novel olfactory-cued navigation task (Fig. 1a-c). Animals had to navigate to cued goals within an environment that was reconfigured multiple times within the same session; animals had to learn and re-learn discrete spatial contingencies in blocks of trials. Each trial in the task began with the animal sampling an odor at the central start location. The particular odor the animal was presented with indicated that there was a reward available at one of four remote goal locations. Animals learned this odor goal association over the shaping period, and were able to go to the correct goal with high accuracy (Fig. 1f), indicating a high confidence with the basic task structure. In blocks of trials within a single session, the location of a barrier within the maze was moved to one of three discrete locations (Fig. 1a). The location of the barrier determined the correct route to each goal (Fig. 1b, Fig. 2b).

**Figure 1.**
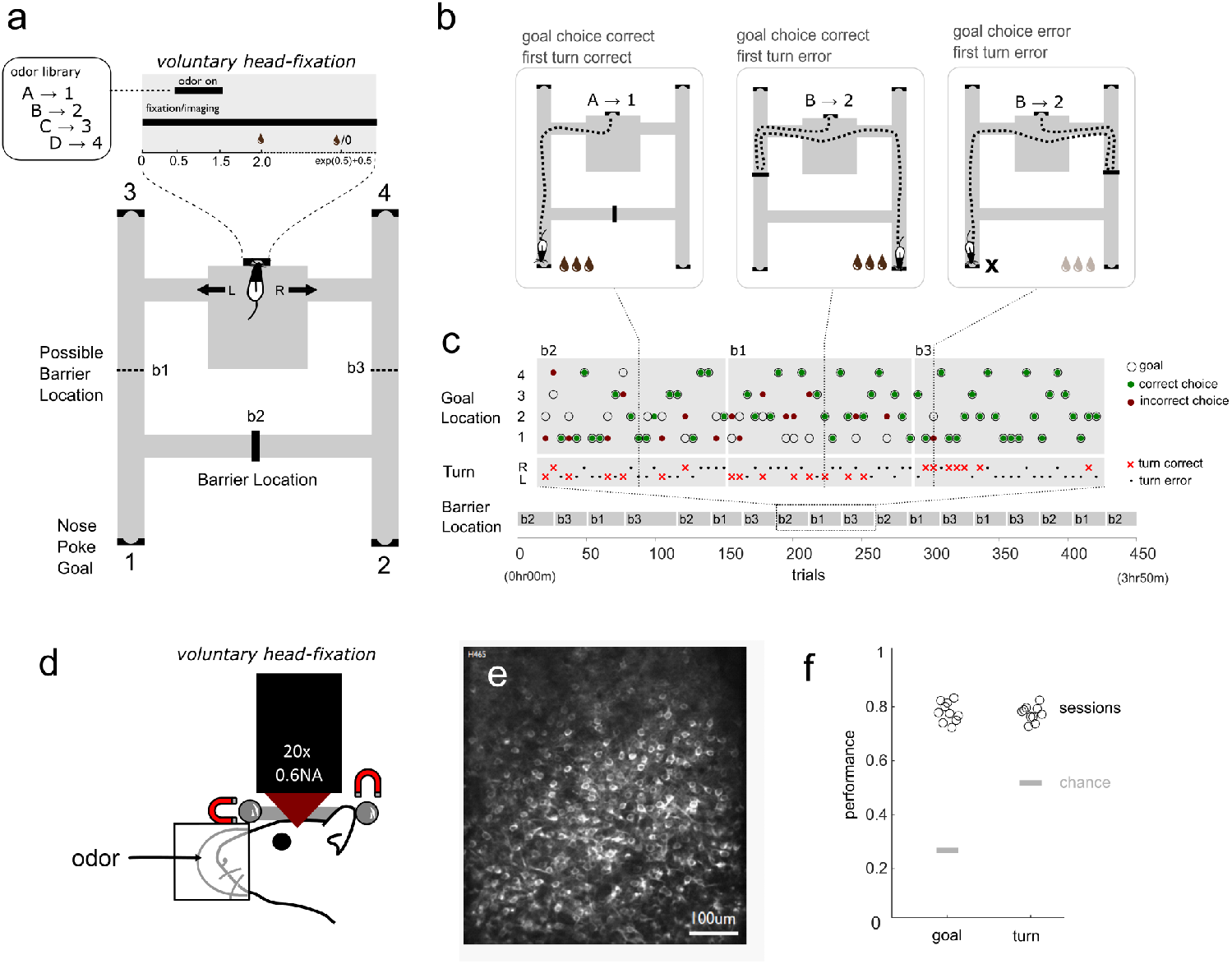
Rats perform a flexible navigation task. **a** – Animals perform the flexible navigation task in a maze. In the central chamber, animals are presented with an odor during a voluntary head-fixation period. Following the head-fixation animals choose to turn left or right to proceed to one of four remote goal locations. The presented odor indicates which goal location has a reward available. Animals have to make an immediate left or right turn decision to navigate to the correct goal. Within the maze, there is a single barrier in one of three locations which is moved in blocks of trials within the session. **b** – Showing three example trials from a session. Animals can make a correct or incorrect goal choice and the first turn the animal makes can either be correct or an error. **c** – The barrier location is changed within blocks of trials within a single session, the trials for three example blocks are shown. **d** – Voluntary head-fixation during odor presentation, the animal is trained to maintain its head in precisely the same position, aided by a light magnetic force and bearing system. Within trial stability and across trial repeatability allows 2-photon calcium imaging of neurons in hippocampal region CA1. **e** – A subset of cells in CA1 express the calcium sensor GCaMP6f, which allows the monitoring of the activity of a large number of neurons. **f** – Over all sessions, performance is generally high for both the goal choice (chance is 0.25) and the first turn (chance is 0.5).

**Figure 2.**
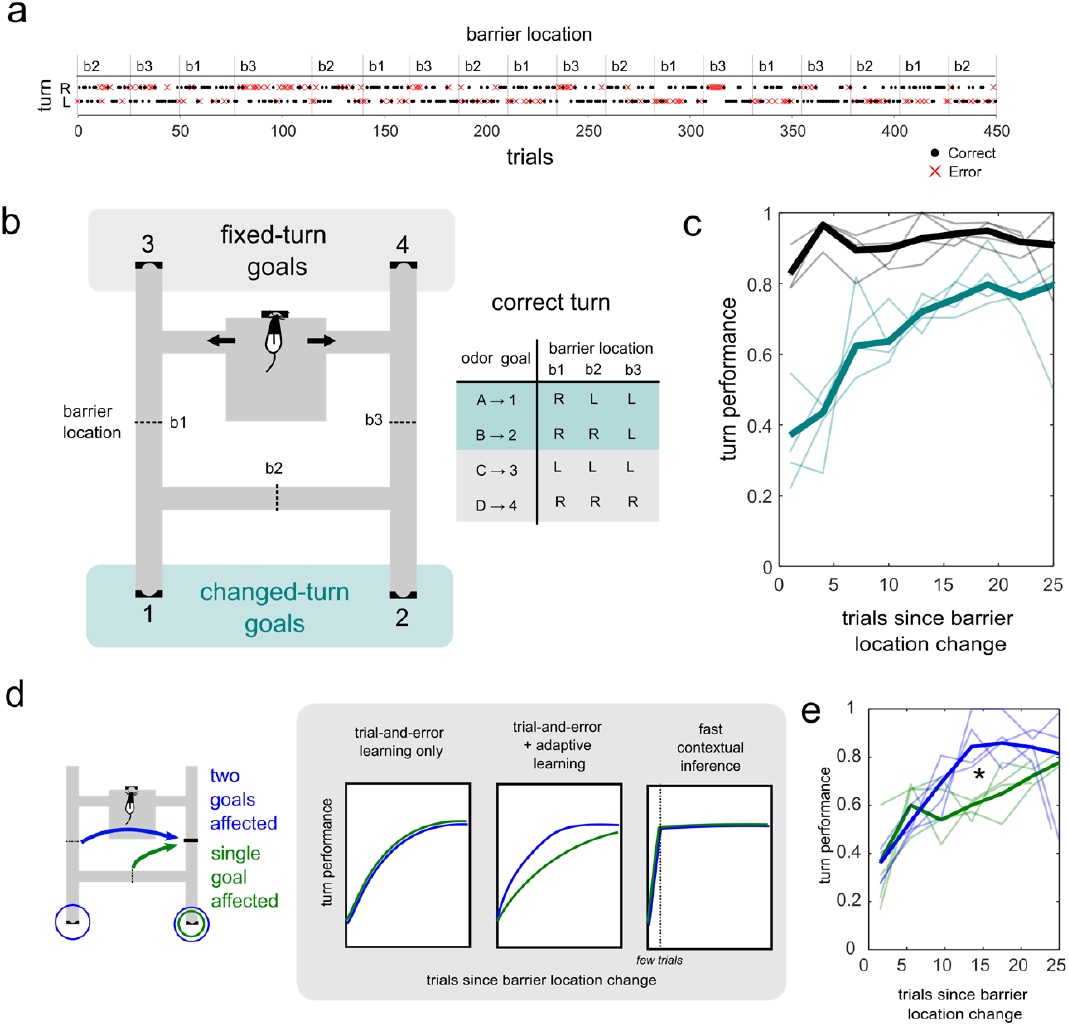
Rats learn new routes in a changing environment. **a** – A full session (same as shown in Fig1) showing turn decisions and the blocks of trials of the different barrier location. **b** – The goal locations in the maze could be characterized as *changed-turn goals* when the correct turn response was changed in different blocks of barrier location, and *fixed-turn goals* when the correct turn response was constant across all barrier locations. **c** – Learning curves for the two goal types, in bold shows the average learning curve and thin lines show individual animals. **d** – Diagram of two block transition types and predicted learning curves. In the diagram the colored arrows represent the movement of the barrier from one location to another. For the blue transition, the correct turn for two goal locations (circled) are affected. For the green transition one a single goal location (circled) is affected. The panels illustrate the predicted learning curves for three types of behavioral strategy. **e** – Learning curves for the two of two block transition types in bold shows the average learning curve and thin lines show individual animals. * – significant difference between the two block transition types (β_ntrial:blockType_ = 2.04, p = 2.08×10^−6^).

### Animals learn new routes in a continual spatial learning task

For two of the goal locations, the correct turn response must be *changed* after certain block switches; for the other two goal locations, the turn response is always *fixed* regardless of what position the barrier is in (Fig. 2b). We looked at the profile of performance for the first turn following a block transition; performance improved as a function of distance from the block transition for changed-turn goals (Fig. 2c) indicating that animals learned the new correct first turn that corresponded to a new route for a given goal. In contrast for fixed-turn goals, performance remained high, indicating animals were able to maintain a fixed policy for some conditions despite the environmental changes going on.

In principle, animals could use a contextual switching strategy to solve this task; the same conditions are revisited multiple times every session, so the animal would have to retrieve the correct set of actions previously learned for each of the three conditions or contexts. Such a strategy would predict fast switching of behavior following a block transition, for instance using state inference (Gershman et al., 2014; Wilson et al., 2014), but this expectation was at odds with the observed gradual learning curves (Fig. 2d,e).

Such gradual or incremental learning is more typically associated with trial-and-error learning, a strategy where an agent learns about the correct action separately for each stimulus (in this case, the odor-goal pair) based on the direct outcomes of its action (Daw et al., 2011). In its simplest form, this strategy would predict learning rates that depended only on the direct experience animals had with each odor, which is the same for all types of blocks. However, animals showed different profiles of learning in different block transition types, in deviation from this simple view (Fig. 2d,e). Specifically, we observed faster learning in block transitions where more than one odor/goal had a changed route (Fig. 2e), indicating that learning was in some way adaptively modulated based on changes in the environment. We quantified these effects using a logistic regression to analyze animals’ turn choice following a block switch for changed goals (Fig. 2e). We found the number of trials following a block switch was a significant predictor of the correct turn (β_ntrials_ = 2.60, p = 2.11×10^−10^), confirming that animals were learning the correct action, and that this learning significantly interacted with the block type (one vs. two goals changed routes, β_ntrial:blockType_ = 2.04, p = 2.08×10^−6^), indicating that animals are sensitive to broader environmental variability to adjust their learning strategy.

### Neural responses in the hippocampus encode odor but not upcoming turn or barrier location

During the odor presentation, animals had been trained to voluntarily head-fixed beneath a two-photon microscope (Fig. 1), which, combined with an optical cannula and transgenic expression of GCaMP6f in pyramidal neurons, allowed us to monitor the population dynamics of a large number of dorsal CA1 neurons (n = 1143 over 10 sessions, session length 3.9 ± 0.8hrs mean ± std). The voluntary head-fixation system consists of a magnetic bearing system that aligns the head to the same position with micron-scale repeatability from trial-to-trial and animals can engage and disengage entirely at will (Rich et al., 2024). Voluntary head-fixation provides stability to perform population calcium imaging and allows us to unambiguously attribute any differences in neural signal in the hippocampus to cognitive rather than positional determinants (Liberti et al., 2022).

The critical decision-making during this task is sequestered within the epoch when the odor is presented and the animal has to make an immediate navigational decision (turn left or turn right, Fig. 1a). This decision is based on the integration of two distinct sources of information: the odor that is presented and the location of the barrier in the maze. We performed a linear decoding analysis to detect which of three behavioral variables: odor, turn, and barrier location were present in the ensemble activity. As expected (Igarashi et al., 2014), odor identity was decodable from the activity of cells in each trial shortly after, but not before odor presentation (all sessions p < 0.01, Fig. 3b). There was no pattern of significant decoding of the upcoming turn the animal was about to make (almost all time points for all sessions p > 0.01 Fig. 3b).

**Figure 3.**
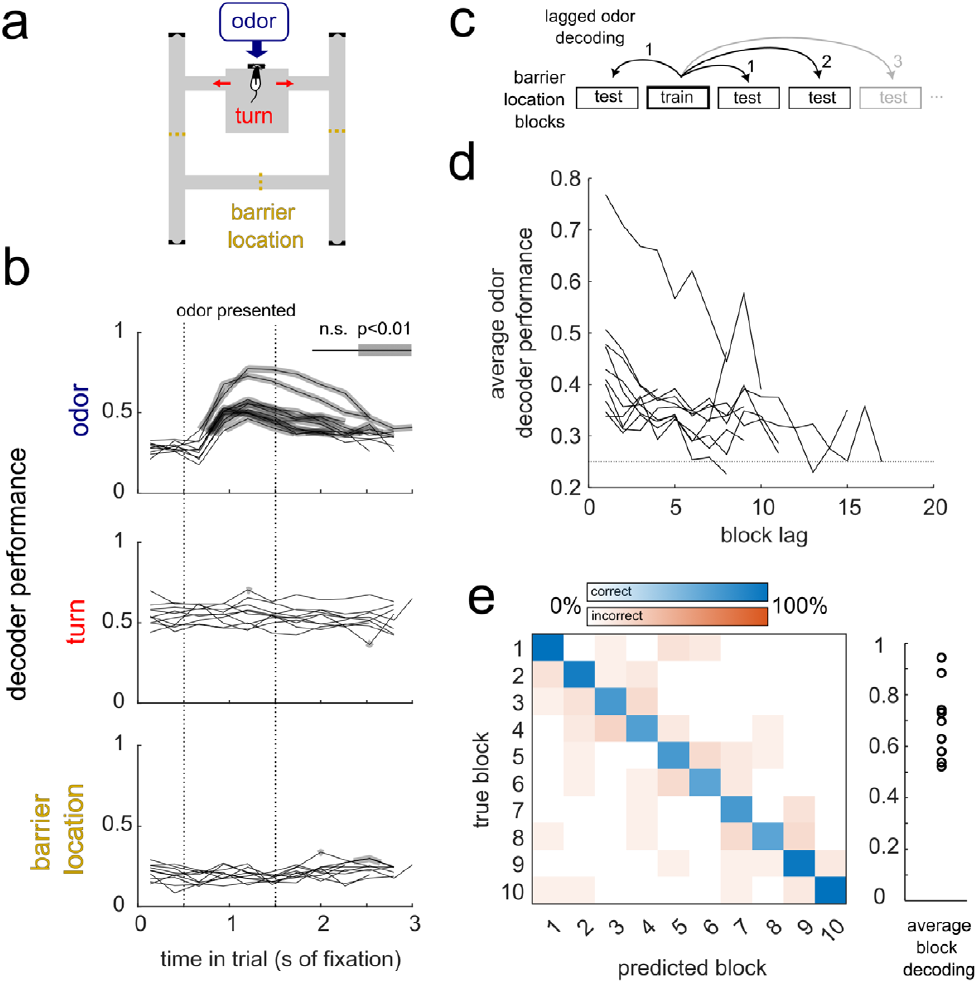
Decoding of odor, but not other task variables. **a** – Illustration of the three factors in the task: The odor presented to the animal, the turn the animal takes and the location of the Barrier. **b** – The time courses of decoding performance during the head-fixation (recording) period in the task. Each line is an individual session. The lines are thickened where the decoding performance exceeded the p = 0.01 proportion of the null distribution, indicating above chance decoding performance. **c** – illustration of the lagged block odor decoder cross-validation scheme. **d** - The average performance of a whole-trial odor classifier as a function of the lag between training and testing. The lines show individual sessions; the nominal chance decoding is 0.25. **e** – Block decoding confusion matric for a single session. Plot on the right shows the average block decoding across different sessions.

To ask whether the barrier location was represented in the ensemble, we used a decoder that was trained and tested over different blocks within the session (this was to ensure that we would be detecting consistent representation of barrier location - see methods). We were unable to decode the current barrier location when a classifier was trained on other blocks (almost all time points for all sessions p > 0.01, Fig. 3b), indicating that there was no reflection of the barrier location context in the hippocampal code during the decision-making period. More sophisticated classifiers that could account for a structured drift of context, that for instance preserved some consistent geometry between conditions (Keinath et al., 2022) also did not reveal any contextual coding (Fig. S1).

We next investigated whether the representation was stable over the multi-hour sessions. We hypothesized that a dynamic representation may facilitate some aspects of the behavior we saw; specifically we wanted to test whether ongoing learning might be facilitated by representational drift (Rule et al., 2019), which would imply an ongoing change in neural activity through the session. We trained odor classifiers on single blocks of trials, and evaluated them for other blocks (Fig. 3c). Looking at the average performance as a function of the lag between the training and the test blocks, we found that there was a gradual decline in the decoder performance for more distant lags. Additionally, the block identity itself (rather than the barrier location) could be decoded, indicating a robust implicit encoding of time within the session (Yang et al., 2024).

### Representation change is apparent at the individual and ensemble level and is continual throughout the session

Inspection of individual units revealed clear non-stationarities in the responses of cells across the recording sessions. Neurons showed a long-tail distribution of the number of trials activity, and each trial showed a sparse recruitment of ∼5% of neurons (Fig. S2). We observed a large variety of individual cell responses, with cells both turning on and off during a session, and with both gradual and abrupt time courses (Fig. 4a-c).

**Figure 4.**
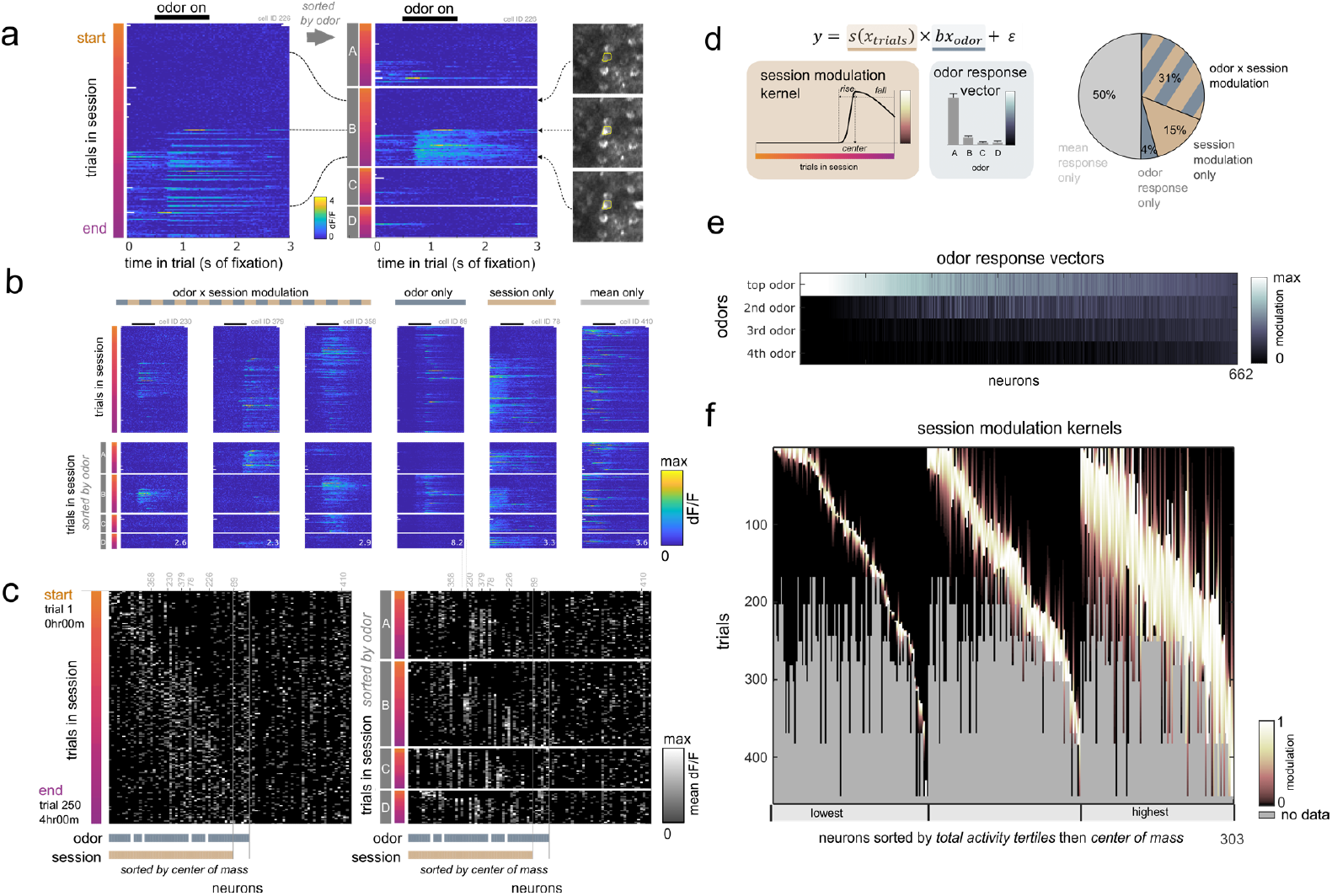
Neurons respond to odor and modulated within the session. **a** – An example neuron showing the individual trial fluorescence activity within the head-fixation period. The first plot is sorted in order of the trials within a behavioral session, and the second plot is the same data, sorted first by the odor that was presented to the animal that trial. The periods of time that the odor was presented within the trial is shown as a black bar on the top of the plot. The three subpanels on the far right show the averaged image centered around that neuron for three selected trials throughout the session, with the dotted lines following the trials back to the session ordered plot. **b** – Neurons from the same session showing examples of the types of odor and session modulation that was observed. Each column is a single neuron, with the two orderings for each neuron as in A. The peak dF/F values are shown at the lower right corner of each plot. **c** – The average activity, following odor presentation, for each trial for all neurons that passed the minimum activity threshold within a single session. Each column is a single neuron, and the color scale corresponds to the average dF/F on each trial, following odor presentation. Before averaging, transients were detected and any non-transients in individual trials are set to zero. The neurons are sorted in three blocks, session modulated neurons, odor only modulated neurons and mean response only neurons. Session modulated cells are sorted within the category according to the center of mass of the session response kernel. **d** – The statistical model used to capture the average responses of individual cells (y) as a product of a parametric session response kernel and a characteristic odor response vector. To the right is the proportion of neurons in each category, showing a significant contribution of one or both of the two factors. **e** – The distribution of the normalized odor responses for all neurons. Neurons are sorted by the lifetime sparseness with respect to the estimated odor vector, and the odors are ordered for each neuron according to their responsiveness. **f** – Showing all session modulation kernels for all neurons that showed a significant modulation across the session. Each column is an individual neuron. The modulation kernels are aligned to the first recorded trial in the session, and are only shown for the number of trials within each session. The neurons are sorted first into total activity terciles according to cumulative sum of the session modulation kernel. Within each tercile neurons are sorted by the center of mass of the kernels.

As well as strong responses to single odors, we noticed that cells also responded to multiple odors at varying levels, while being modulated by the overall time within a session. We used an encoding model for the individual neurons that explained the average activity of a cell within a trial as a multiplication of a parametric temporal response kernel over a session and a characteristic odor response vector (Fig. 4d). This approach allowed us to test whether a cell was modulated by time during the session, by the presented odor, or by a multiplicative combination of the two (Fig. 4d). We found cells in all categories, with a large proportion of cells showing this conjunctive session and odor coding. We also found a distribution of the odor responsiveness of cells, with a large proportion of cells showing a strong response to a single odor (Fig. 4e). Inspection of the significant session modulation kernels revealed a broad distribution of shape profiles for cells, with some cells showing small firing fields over the session’s timeline, and others broader activation (Fig. 4f); these responses tiled the session (Fig. 4c, f). There was no correlation between the strength of the session modulation and the isolation quality of the ROI (Fig. S3) indicating that the modulation we saw was not a result of movement of the imaging plane (Fig. 4a). Finally, we noticed that many of the cells had a large transient early, or on the very first trial of their transition from silent to active (Fig 4a, Fig. S4). These large transients are likely plateau potentials that are the initiation events for behavioral timescale plasticity (Bittner et al., 2017).

### Representational change increases with learning

We next asked whether the representational changes we observed were modulated by any behavioral features related to learning. One possibility is that representational change is entirely dependent on time, encoding the elapsed time within a session at a constant rate. Alternatively, there may be factors in the experiment that are related to learning that show correspondence with the representational change. To begin, we calculated a pairwise population vector correlation matrix to visualize the change in the ensemble response over a session (representational similarity matrix Fig. 5a) and plotted the average neural similarity as a function of trial lag (Fig. 5b). In line with our univariate description and previous work (Manns et al., 2007; Yang et al., 2024), we saw that time within a session was a strong predictor of neural similarity, evidenced by a gradual fall off of neural similarity with trial lag. As expected from the decoding analysis and univariate encoding analysis, pairs of trials with the same odor were more similar than those with different odors (Fig. 5b). Breaking the category of same odor trial-pairs down further, we found that the average population vector distance was systematically increased for changed-turn odor/goals compared to fixed-turn odor/goals (Fig. 5b,c; see Fig 2b), suggesting that the learning demands of the task increased the rate of change of the population.

**Figure 5.**
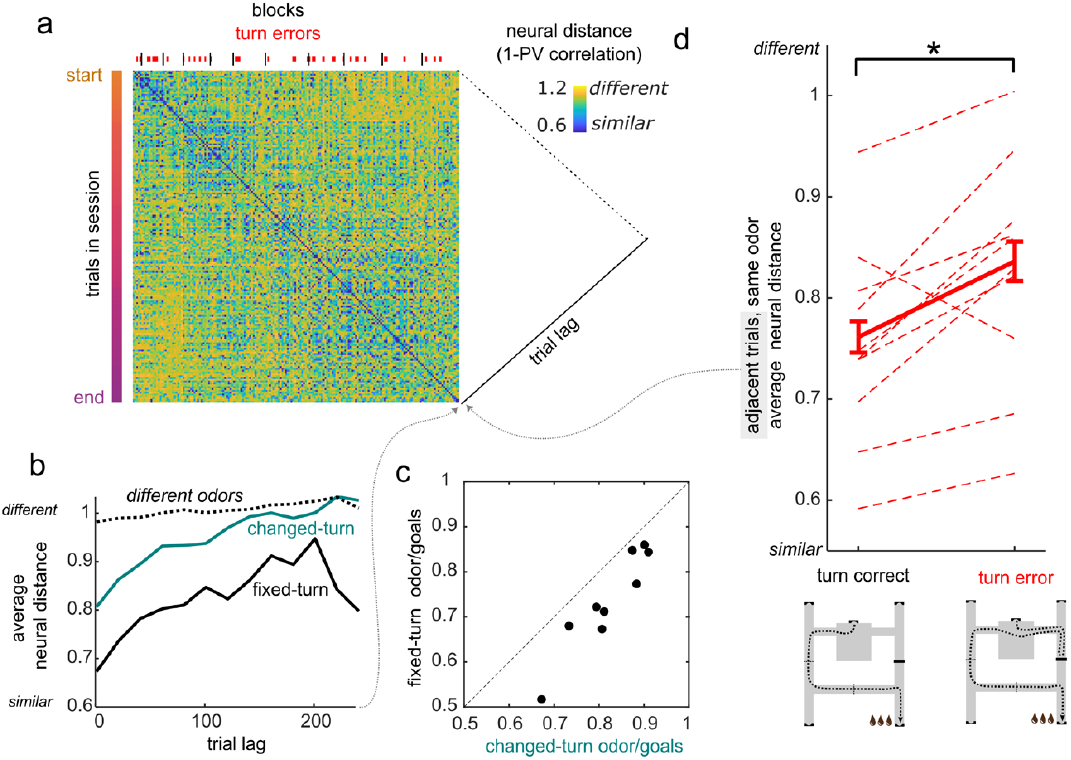
Neural change is modulated by factors of learning. **a** – Representational similarity matrix for a single session, each element is the trial-to-trial similarity, the neural distance is 1 minus the population vector correlation. Along the top edge shows the blocks and the turn errors within a session; along the side edge shows the trials in the session (matrix is sorted by time) **b** – the representational similarity matrix is collapsed along the diagonal to generate the similarity with respect to trial lag for a single session. The three conditions correspond to three trial-to-trial comparison types. Solid lines are between trials with the same odor, in the two categories (Fig. 2), dotted line is between trials of different odors (bin width 20 trials). **c** – The average distance between pairs of trials of the same odor for fixed vs changed turn conditions for the first trial lag bin in b (1-20 trials), shown for each session separately. **d** – The average neural distance between adjacent trials of the same odor, which is the collapse of the representational similarity matrix along the off diagonal, separated as to whether the animal made a turn error on the first trial. Dotted line represents the average values for individual sessions. * – p = 0.005 for main effect of error in mixed-effect model.

To investigate the fine time-scale changes in neural representations and to test them statistically, we examined neural representations between adjacent trials (consecutive trials in the experimental sequence). This approach allowed us to investigate how representational changes might be related to learning on a trial-by-trial level. We found that there was a significant effect of the fixed-turn category on the neural representational distance between adjacent trials (β_fixed_ = -0.09, p = 0.016) even accounting for the absolute time difference between trials (β_timeDiff_ = 0.00, p = 0.169). Looking at adjacent trials also allowed us to exclude the possibility that the difference between fixed-turn and changed-turn odor/goal trials was driven by exposure frequency (fixed-turn odor/goal trials were presented less frequently than changed-turn trials).

In many computational models of learning, learning is driven by errors, which can represent a discrepancy between the predicted and actual outcome. We asked if the neural representational similarity between adjacent trials for the same odor for changed-turn odor/goals was modulated by turn errors. We found that neural distance was increased between adjacent trials of the same (changed-turn) odor if there was an intervening turn error (i.e. an error on the first trial, Fig. 5d, β_error_ = 0.07 p =0.005), this difference was not explained by absolute time difference between trials (β_timeDiff_ = 0.00, p = 0.46). This strongly indicates that the process of representational drift serves a purpose in the ongoing learning process.

### Representational change enables flexible re-learning without forgetting

In order to understand the mechanism of how dynamic, non-stationary neural data could give rise to the patterns of learning we observed in the behavior, we constructed a simple feedforward neural network model. The input layer corresponds to the CA1 response to a given odor, and the output units model the decision of whether to turn left or right, for instance in a downstream target (Fig. 6a). A weight matrix representing synaptic weights is updated based on a supervised Hebbian learning rule. Such learning rules typically rely on decay, normalization or depression of weights to achieve re-learning by erasing old weights; for instance, if the contingencies for a certain stimulus are reversed. However, the erasure of previous associations following re-learning is at odds with experimental evidence that associations can be recalled later following reactivation of tagged engram cells (Liu et al., 2012). To account for this, we included a constraint that a synapse could only *increase* in weight and would not decay or be depressed, allowing the possibility for associations that could persist in time. Under this constraint, a static input representation of the odor stimuli prohibits any re-learning since the previous association would compete with any new one (Fig. S5). We reasoned that drift in the input representation would allow re-learning by 1) moving activated inputs away from the previous rules in the weights and 2) providing a fresh, un-associated set of weights on which to make new associations.

**Figure 6.**
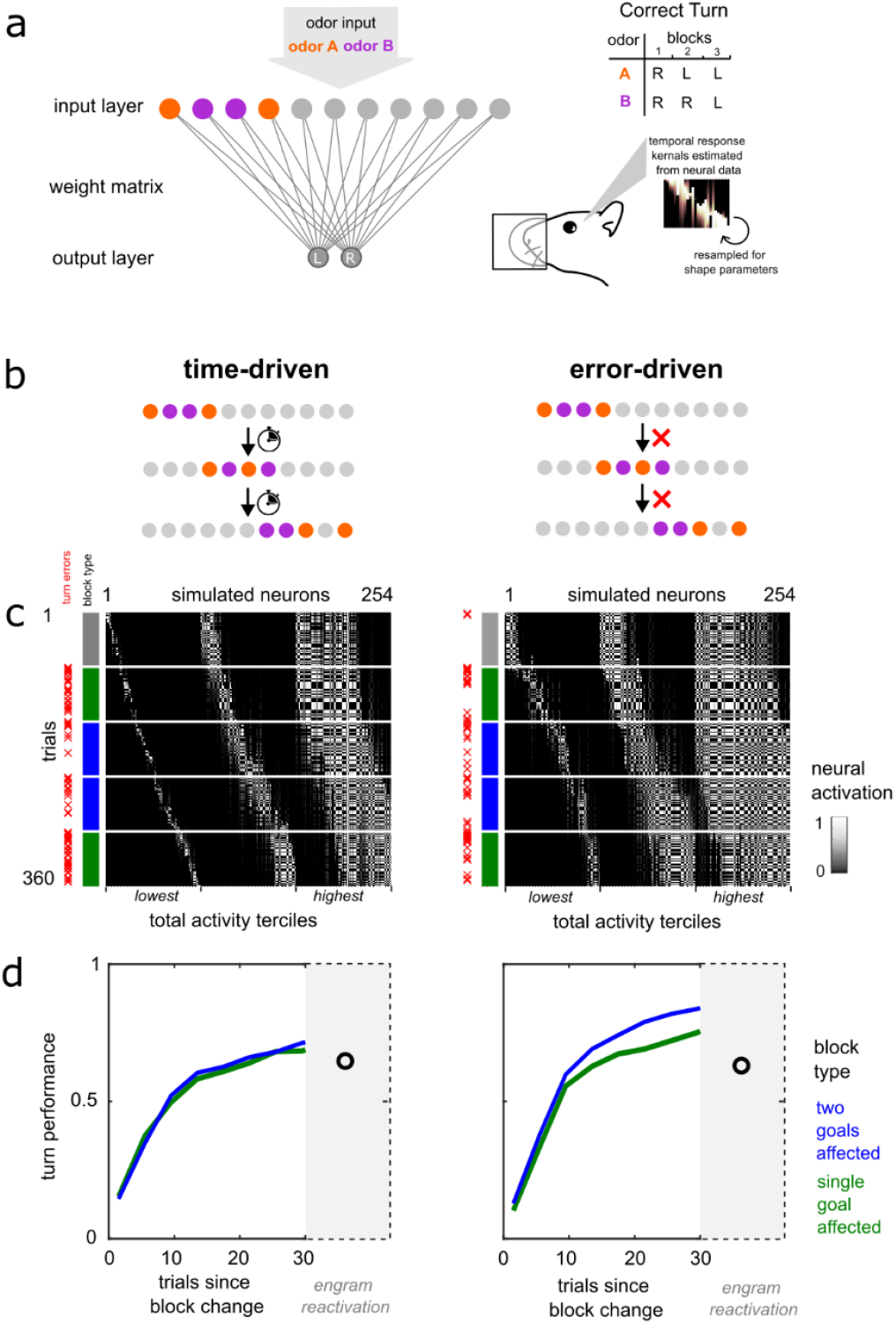
Error-driven change enables adaptive re-learning. **a** – Feedforward neural network architecture. The Input layer is activated by the presentation of two odor, which project to an output layer with a turn left and turn right unit. The network had to perform a simplified version of the behavioral task, using only the two changed turn odors. The coarse temporal/session modulation of simulated cells is determined by resampling the session response kernels estimated from the neural data. **b –** The fine time-scale of the session modulation was tested in two conditions i) time driven which advanced the representation on every trial or ii) error-driven which advanced the representation following errors **c-** Activity of simulation neurons across the session. Neurons are sorted in the same manner as Fig. 4f, first into 3 terciles based on total activity and then, within each quartile by the center of mass within the session. **d** – The average learning curves of the network as a function of the number of trials since the block change (barrier changes location). The categories are the same as for Fig. 2b,c: block transitions with two goals affected and block transitions with only a single goal affected. The last data point is the performance of the network during an engram reactivation test, where activations from previous points in the session are sent through the final weight matrix that has been learned at the end of the session.

We imposed representational change in the input layer in our network to mirror the neural data that we recorded in CA1 and investigated two conditions for how the drift/change was driven (Fig. 6b). In the first condition, the representational change occurred as a fixed amount per trial (time-driven), and in the second change only occurred in the input layer when the agent made a turn error (error-driven) - more closely mirroring our observation from the neural data. In both cases, the network was able to both re-learn new rules following block transitions (Fig. 6d). The drift of the input vector opened up a fresh, unused set of weights on which new association could be made, and the previously learned set of weights for changed units would not contribute to the action since the input units were now silent (Fig. S5).

Only the error-driven change model exhibited the learning rate difference between the different categories of block transition that was seen in the behavioral data (Fig. 6d *c*.*f*. Fig. 2e). In the time-driven drift, the rate of change was constant despite ongoing changes in performance. However, when the drift was driven by turn errors, this caused an increased level of drift in blocks with more errors (blue, two goal changed condition). Since the drift affects the representation generally, for both odors, this is then seen as an increase in the overall learning rate. After an error for one odor there is a weakening of the rule for the other odor; when both odors need to be relearned, this is beneficial, as it weakens the previous, now incorrect association for the other odor. In the case where only one odor is relearned, there are fewer errors and the generalized unlearning is balanced by the ongoing associative strengthening that occurs on correct trials.

Finally, at the end of a simulated session, following learning, we simulated a hypothetical engram reactivation experiment; we reactivated the patterns of input from pervious trials in the session and presented them to the final end-of-session synaptic weights. The network was able to retrieve the correct rule better than chance, indicating that a record of the previously appropriate actions was successfully stored in the final weight matrix (Fig. 6d, Fig. S5). Although we used the full distribution of time scales as estimated from the neural data, investigation of the effective weight matrices (Fig S – effective weights) indicated that only the fast-modulated simulated neurons were contributing to this recall.

## Discussion

We present a study to examine the neural bases of how animals adapt and learn in changing environments. We designed a novel learning paradigm that challenged animals to learn and re-learn, revisiting repeated conditions, throughout long, multi-hour sessions. We used magnetic voluntary head-fixation during the crucial decision-making epoch in the task, allowing us to monitor large numbers of pyramidal neurons in the CA1 of dorsal hippocampus. We did not observe a representation of the maze configuration in the neural data, nor behaviorally the sort of fast rule switching that such a representation could support. Rather, the neural task representation, which primarily encoded the instructional odor stimuli, changed steadily throughout the session. On a trial-by-trial level the representational change was increased following errors, indicated the role of the change for ongoing learning in the task. Although often considered a problem for the stable readout of activity in downstream circuits (Rule et al., 2019), we show how this representational change may in fact serve a useful purpose by allowing flexible behavior, enabling the learning of new rules and the suppression or extinction of previous ones. This architecture resolves a tension between stable memory storage and representational drift, as previous associations are stored and may be recalled from latent synaptic weights. Finally, the modulation of the rate of drift by errors emerges as a mechanism by which learning rates can be altered in environmental conditions with varying volatility, indicating a role for drift beyond simple temporal encoding.

### Drift in Hippocampal representations

Activity in the hippocampus delineates spatial (Leutgeb et al., 2005; Muller and Kubie, 1987) latent (Pastalkova et al., 2008) and abstract (Nieh et al., 2021) contexts. Reactivation studies have shown that this activity confers the contextual specificity of long-term memories (Liu et al., 2012). However, hippocampal activity is increasingly appreciated to change over time, showing different responses even when identical environmental conditions are experienced (Bladon et al., 2019; Kentros et al., 2004; Mankin et al., 2012; Manns et al., 2007; Ziv et al., 2013). In our work, we make a number of contributions to the understanding of representational drift and representations in the hippocampus.

Much of the previous information about hippocampal drift comes from looking at representations across sessions or days (Mankin et al., 2012; Ziv et al., 2013), although some studies have looked at fast time-scale changes (Bittner et al., 2017; Monaco et al., 2014). The long sessions we used provided the temporal sampling required to estimate fast and slow changes in coding. We found a diversity of time courses of modulation of individual cells,

The hippocampus also encodes non-spatial elements of the environment (Wood et al., 1999; Manns et al., 2007; Igarashi et al., 2014), we demonstrate that representational changes are not exclusive to the spatial representations in the hippocampus, which have been the focus of previous investigations of drift (Kentros et al., 2004; Geva et al., 2023; Khatib et al., 2023). Head-fixation allows an unambiguous physiological demonstration of drift since potential positional determinants of neural coding are eliminated (Liberti et al., 2022); two-photon calcium imaging is ideally suited for the study of representational drift (Driscoll et al., 2017) by eliminating any concerns that change is due to electrode movement.

### State representations and mechanisms of learning

Learning can be envisaged as the process of forming synaptic links from the representation of the current state to the appropriate action (Brown and Sharp, 1995). When the environment changes, the appropriate actions may also change. However, the encoding of new information can lead to catastrophic interference (McCloskey and Cohen, 1989) a well-known problem facing a neural network when existing information is overwritten by the new. We have demonstrated how a dynamic representation can ameliorate this concern and provides a scalable solution for continual learning (Driscoll et al., 2022).

In our model of drift and learning, the dynamic state representation serves two roles when contingencies change. The first is to suppress the expression of previous learned information, which is stored in the downstream weights of now silent neurons, which represents an orthogonal neural population. The second is to provide a new set of active neurons with fresh downstream weights for new information to be stored. This idea has been proposed as a mechanism for memory encoding (Mau et al., 2020; Driscoll et al., 2022; Antony et al., 2024) that would involve a subset of neurons that are recruited to be part of the memory trace (Zhou et al., 2009).

Perhaps the closest previous theoretical model to ours is Razmi and Nassar’s (2022) proposal of a dynamic state mechanism underlying adaptive learning. Our data provide direct cellular support for this concept and connect it to phenomena of hippocampal representational drift. The prominent temporal context model (Howard and Kahana, 2002) proposes that a gradual drift of a contextual memory representation explains contiguity effects in memory recall. This line of work emphasizes the role of experiential effects (stimulus encounters rather than the passage of time per se) in driving such change, and evidence from both animals (Geva et al., 2023; Khatib et al., 2023) and humans (DuBrow et al., 2017) has suggested that the rate of drift in the hippocampus is not solely temporally driven, but is influenced by experience and behavior.

### No upcoming turn activity seen in the hippocampus

We did not see any decoding of the turn the animal made in the neural activity in CA1. This was somewhat surprising given the existence of prospective trajectory modulation of place activity in the hippocampus (Frank et al., 2000; Wood et al., 2000); however such “splitter cell” activity is not always observed (Berke et al., 2009; Duvelle et al., 2023). Furthermore, most studies observe prospective activity during locomotion either on a track or a running wheel (Duvelle et al., 2023), indeed in a brain-wide survey during a head-fixed visual decision task, the hippocampus was one of the few areas that did not show upcoming choice coding (Steinmetz et al., 2019). Many studies that observe trajectory-modulated activity have the structure that memory of the prior trial uniquely determines the upcoming response, whereas in our task the decision to turn left or right is more complex, requiring the integration of information about both the current odor stimulus and the previous history.

### Absence of latent context representation

We did not observe any reflection of the latent context, the closed barrier location, in our neural data; our analysis would have detected invariant stationary code as well as a drifting code with consistent geometry or similarity for contexts (Keinath et al., 2022). Although well-trained, the animals in this study did not immediately deploy the correct rules in a given context block despite having experienced it many times before. Latent context models (Gershman and Niv, 2010; Wilson et al., 2014) suggest that previously learned rules can be accessed through inference of a latent context representation, providing behavioral benefits such as fast learning or generalization. The lack of contextual reactivation in the neural data during our task is therefore, entirely consistent with the lack of fast contextual switching in the behavior.

A caveat to our findings is that we are not recording from the hippocampus throughout the whole task. Although we designed the task to sequester the crucial decision-making period in the head-fixation imaging epoch, it may be that a latent context representation would be activated at other timepoints, for instance when the animal is actively navigating. However, even with continuous recording, it may prove difficult to definitely interpret any putative contextual signal since there would be covariate positional changes such as slowing down before encountering a closed barrier.

Contextual coding in the hippocampus is frequently observed when context is overt, either spatial context (Muller and Kubie, 1987,Kubie et al., 2020) or stimulus-defined contexts (Markus et al., 1995; McKenzie et al., 2014). However, other studies with a similar, block-based switching task to ours did not any behavioral context switching in rats or any reflective contextual neural representation in CA1 (Bladon et al., 2019), suggesting that the reliance on incremental learning over contextual inference may be a common strategy adopted by animals in certain conditions. Simple incremental learning algorithms fail to account for learning phenomena such as savings over repeated reversals and reinstatement following extinction (Redish et al., 2009) where animals show patterns of learning that deviate from what would be expected for a direct incremental learner. In our study, we saw other deviations from simple incremental learning (Fig. 2e) that can be accounted for by the effective learning rate being modulated by errors.

### Incremental learning and state inference

Trial-and-error incremental learning and latent state inference have been viewed as distinct cognitive processes with specialized neural mechanisms that can both contribute to behavior (Bouchacourt et al., 2022). In our account, these two seemingly disparate processes can in fact be understood as separate operations of the same system (Yu et al., 2021). A dynamic state representation provides a fresh neural substrate for continual learning of new information and previously learned information is stored and can be recalled via state inference. Mechanistic questions that have risen from these separate accounts can be reinterpreted in terms of the control of hippocampal representational dynamics. The rate of change of the state representation may be used to modulate learning rates (Razmi and Nassar, 2022), and the error-driven change we observe is a simple implementation of what might otherwise be a complex estimation of environment volatility (Piray and Daw, 2024).

What factors might influence reliance on contextual or trial-and-error incremental learning strategy? The frontal cortex is thought to be critical for latent state inference (Wilson et al., 2014) which could act as a controller to reactivate previously active patterns in the hippocampus and recall the information. While a static context representation allows rapid access to a fixed set of rules, such a system may take time to learn and potentially less useful in dynamic environments, especially where there is uncertainty about the statistics of change.

Although in our experiment we don’t see any instantiation of the latent context, our model predicts that reactivation of patterns of previous activity from a previous block, for instance cell-specific optogenetic stimulation (Rickgauer et al., 2014; Robinson et al., 2020), would cause the animal to retrieve the appropriate rule from that block, rather than simply providing the percept of the stimulus. Furthermore, we would predict that in conditions where fast contextual switching was seen in the behavior, a concordant contextual representation would be seen in the hippocampus.

The distribution of modulation rates across the population we observe in our data also suggests a reservoir of temporal scales that would be useful for learning associations across different time scales. We found that only the fast-modulated neurons would be contributing to recall of the block specific information since it was varying at a similar timescale. In the same way that place cell field propensity is broadly distributed across the population to enable efficient coding of large and small environments (Eliav et al., 2021; Rich et al., 2014), a multi-scale distribution of temporal modulation would allow a similar advantage for making associations over various times-scales.

#### Outlook and Future Directions

Representational drift has also been observed in other brain regions (Deitch et al., 2021; Driscoll et al., 2017; Karlsson et al., 2012; Schoonover et al., 2021; Tsao et al., 2018) indicating that it is a general feature of many neural systems. Understanding the purpose and mechanisms of dynamic, non-stationary neural circuits generally is a pressing problem in contemporary neuroscience (Driscoll et al., 2022; Rule et al., 2019) and advances in longitudinal neural recording capabilities (Rich et al., 2024) have created an opportunity for the continued investigation of this phenomenon.

## Acknowledgements

This work was supported by grants from the Simons Collaboration on the Global Brain (D.W.T.), NIMH R01MH136875 (N.D.D.) and ARO W911NF2410183 (P.D.R.). We thank the Tank and Daw Labs for valuable discussions, especially J. Chang, E. Diamanti. We thank S. Koay for cell extraction software and assistance.

## Data and Code Availability Statement

Upon publication, data will be made available on a publicly accessible data sharing site. Code for analyzing the data will be available on GitHub

## Inclusion and Ethics Statement

All animal procedures were performed in accordance with the Princeton Universty Institutional Animal Care and Use Committee (IACUC) guidelines and approved under protocol #1837

## METHODS

### Animals and Surgery description

All procedures performed in this study were approved by the Institutional Animal Care and Use Committee at Princeton University (IACUC protocol number 1837) and were performed in line with the Guide for the Care and Use of Laboratory Animals (National Research Council, 2011). We used 4 male transgenic rats that expresses GCaMP6f under the thy-1 promotor aged between 3 and 9 months at the time of surgery, these rats had were among the animals reported in a previous report (Rich et al., 2024).

Rats underwent surgery to obtain optical access to the dorsal hippocampus. Surgery was performed under aseptic conditions, and animals were anesthetized with isoflurane (3.5% induction, 0.5 -1% for maintenance). Animals received pre-operative buprenorphine (0.02mg/kg) and either post-operative meloxicam (1mg/kg) or buprenorphine (0.02mg/kg) for analgesia. Body temperature was maintained with a heating pad (Harvard Apparatus). The skull was exposed and the periosteum was retracted. A craniotomy was performed on the right hemisphere over the dorsal CA1 region of the hippocampus (4.2AP, 3.0ML) The overlying cortex was aspirated to exposure the fibers of the external capsule, which was carefully retracted to exposure the fibers of the alveus. A thin layer of Kwik-Sil (WPI) was squeezed on the exposed area and the custom conical canula (5/4 mm top/bottom diameter 2.6mm height) was quickly lowered and cemented in place with adhesive cement (C&B Metabond, Parkell). A titanium base head-plate was mounted parallel to the stereotaxic frame and cemented to the skull. The canula was held in place at its rim by three magnets (1/16” diameter, D12-N52, K&J magnets) and was implanted centrally to the head-plate aperture with the cover-glass parallel to the stereotaxic frame. These steps resulted in the cover-glass being parallel and concentric to the head plate for imaging. Following surgery animals were left for ∼1 month before the imaging window was assessed under anesthesia. At this point the kinematic head-plate was installed into the base head-plate.

At the conclusion of the study, animals were anesthetized with an overdose of ketamine xyline and euthanized with a transcardial perfusion of PBS followed by 4% paraformaldehyde. Brains were extracted, and left in fixative for 12 h and then transferred to 30% sucrose in PBS until the brain had sunk. Brains were sectioned on a freezing microtome (Leica) in 50μm sections and floated onto slides. Slides were inspected on a epifluorescence microscope to confirm that the field of view of the cannula was in the CA1 region of the hippocampus as assessed by gross anatomy.

### Imaging system

We performed imaging with a custom two-photon microscope controlled with the Scan Image software (Vidrio) running in MATLAB (Mathworks). Laser illumination was provided by a Ti:Sapphire laser (Chameleon Vision II, Coherent) operating at 920 nm. We used a long working distance air immersion objective lens (20x/0.6NA/13mm WD, Edmunds optics) and a GaAsP photomultiplier tubes (H10770PA-40, Hamamatsu). The beam power was modulated by a Pockels cell (350-80 LA BIC -02 Conoptics) and the power used for imaging, measured at the front of the objective, was 150-200mW. Horizontal scans of the laser were achieved using a resonant galvanometer (Thorlabs). Typical fields of view measured approximately 600 × 600μm and data were acquired at 30 Hz.

### Magnetic voluntary head-fixation

Animals were first trained to voluntarily clip into a magnetic head-fixation system (Rich et al., 2024) that provided micron-scale registration of the head required for trial-to-trial calcium imaging of the hippocampus. Animals were initially trained to do this as previously described (Rich et al., 2024). Briefly, animals were first trained to poke at the stainless-steel nose poke beneath the head fixation system, to receive a chocolate milk drop reward (1 drop, Ensure milk chocolate flavor nutritional shake, Abbott). A variable period was introduced while the animal had to maintain the hold to receive the reward. If the animal made contact with the bearing surfaces, it was given an immediate larger reward (2 drops). The various further steps of contact with the kinematic clamp were shaped using this strategy hold small reward, next step no hold large reward, until the animals were making routine fixations with the magnetic bearing system. At the start of all sessions animals typically were given 10 warm-up trials with shorter fixations.

### Odor-guided navigation task

Once animals had learned to reliably clip-in to the head fixation system they were trained to perform an odor guided navigation task. We delivered odors (acetic acid 20%, 2-propanol, propyl acetate, anethole, trans-cinnamaldehyde, nutmeg, cardamom, clove, rosemary, star anise) to animals using a custom built olfactometer. Air was directed to flow through vials containing odorants using PTFE valves (Neptune Research) at a flow rate of 0.5L/min. At the start of head fixation, blank air was delivered for 0.5s, followed by 1s of odorized air, followed by a scavenger vacuum through the system to evacuate any residual odorant on that trial and for the next trial. Animals were trained that different odors corresponded to different reward location in a maze constructed of enclosed linear track elements constructed of plastic U-channel and acrylic wall and ceiling. Different arms of the maze were made texturally distinct by adding metal elements to the track floor such as small ramps or protruding screw heads. Animals preformed the fixation and odor sampling in a central chamber. Once the animal had made the fixation for the required duration (2-3s), breaking the fixation would open two choice doors (metal duct blast gates; McMaster-Carr) on the left and right side of the animal, actuated with a servo moto. The animal then chose to turn left or right, and to proceed to the remote goal location in the maze. The maze was configured with a single barrier location (metal duct blast gates; McMaster-Carr), that meant that there was a single correct route to each goal, and so a single correct left right initial turn to make after fixation. During navigation the maze room was lit with a dim purple/near UV LED lamp with additional smaller UV LEDs mounted on one wall to aim as an orientation cue. These lights were turned off during imaging to prevent light contamination. The barrier location was changed manually by the experimenter between each block of trials. If the animal nose poked to the correct goal location given the odor presented it received a large reward (5-8 drops). A nose poked at an incorrect location would elicit a short white noise sound and a 30s timeout. During the timeout the choice-doors were closed, preventing re-access to the main chamber. Following a correct goal choice or the end of the timeout, the animal was free to re-enter the main chamber, where poke a rewarded poke at the reset location at the rear of the main chamber, caused the choice doors to close, and the apparatus to be reset for the next trial. Odors were presented pseudo-randomly in block of 12 trials, with 2 trials for each fixed and 4 trials for each changed odor/goal trial types. Blocks for each barrier location were 24 -36 trials long. The task apparatus was controlled using Arduino microcontroller and custom MATLAB software running of a laptop computer.

### Behavioral analysis

We assessed the learning of correct turn using a logistic regression analysis. We only considered trials for goal locations that had changed a route and therefore the correct turn in a new block. The probability of a correct turn was modeled function of the number of trials since the blocks transition and the categorical block transition type. The block transition type was one of two categories that depended on the previous and current location of the barrier. A single goal turn change was a move of the barrier that only affected the correct turn for one goal location and a multiple goal turn change was a move of the barrier that affected two (both) of the (learned) goal locations. The logistic regression included an intercept: to account for the baseline on the first trial of the block; a main effect of the number of trials since the barrier location change, to capture the animals’ improvement over each block; and an interaction term for the number of trials since the barrier location change and the block transition type, to account for differences in the slope the learning curves. We did not include a main effect of the block transition alone, since it would have modeled a difference in performance before the animal knew what block transition type it was in; which was not supported empirically by the data, or in principle since the block transition type order was random. We fit the model across the dataset of animals and sessions using a mixed effects model in MATLAB, including the random effect of animal ID.

### Data pre-processing and neuron identification

To correct for non-rigid brain motion in imaging data, we employed custom MATLAB code based on the NoRMCorre algorithm. (Nieh et al., 2021, Pnevmatikakis and Giovannucci 2017). This method involves dividing the image into overlapping patches and estimating a rigid translation for each patch and frame by aligning it with a template. The resulting translations are then interpolated to form a motion field, which is applied to smaller overlapping patches. The displacement for the x and y axes were determined by the maximum values observed across all patches in each frame.

Fluorescence traces for individual cells were extracted from the motion-corrected images using the Constrained Non-negative Matrix Factorization (CNMF) algorithm (Pnevmatikakis et al., 2016) The initialization of spatial components for CNMF and the classification of these components into cell-like and non-cell-like categories followed previously published methods (Koay et al., 2020) All components were manually reviewed and reclassified if necessary. The ΔF/F for each cell was calculated using the modal fluorescence value from a 3-minute windows as the baseline. Since CNMF only identifies cells with calcium activity during imaging, cells that did not exhibit activity were not included in this analysis. Shape correlation was used as a metric for cell isolation quality. The shape correlation is the correlation of activity in each pixel with the average, reference image for that cell. A change in the shape correlation time course over a session could reflect some change in the measurement, such as a shift in the focal plane. Any units with a clear temporal trend in the shape correlation time course were excluded during manual curation.

We detected transients as periods where the ΔF/F signal exceeded 2 standard deviations in the positive direction for at least 4 frames (133ms). For all analysis we zeroed out all non-transient portions of the individual time-traces for each neuron. Our expression of the GCaMP6f was sparse compared to other systems when contamination of signals from neighboring neurons was not a concern, so false attribution of multiple cells contributing to the same ROI was unlikely. For all analysis, we excluded cells that did have at least one transient in at least 5% of trials within a session.

### Univariate encoding model

The encoding model was formulated as a multiplicative session modulation by odor modulation model. The average calcium response on each trial, following odor delivery, *y* was modeled by a temporal kernel *s* across the trials in a session x_trials_ multiplied with a characteristic odor response vector *b* which has a different value depending on the odor presented that trial x_odor_.

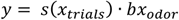

The session kernel *s* was a three-parameter compound gaussian with a center position, and separate rise and fall width parameters. The rise and fall parameters allowed the shape to smoothly transition from a step function to fully flat. The characteristic odor response vector *b* was a positive valued vector. The multiplication of these two terms would give the underlying latent excitability value for each cell per trial. We noticed that residuals of this model fitted to the trial-average calcium data showed strong heteroskedasticity, the variability of the average calcium response increased with the local mean; this is expected since the underlying spiking process is roughly Poisson. We applied the Anscombe transform to the data to stabilize the residual variance, which could then be appropriately modelled with a gaussian distribution for likelihood fitting purposes.

We used the MATLAB function *fminunc* to minimize the negative log likelihood. In order to classify cells, we estimated 5 models for each cell; (1) a mean only model, (2) an odor coding only model, (3) session-time only, (4) the full session-time x odor model and (5) a separate session-time kernel for each odor model.

We used lowest value of the Bayesian information criterion to select the best fitting model for each neurons. We excluded fitted models that showed narrow session kernel(s) width (less than a cumulative sum of the kernel of 6 trial units, which corresponds to a symmetrical gaussian kernel with ∼6 trials s.d.); this constraint corresponded to overfitting of single isolated transients for sparsely firing neurons. Since very few cells (2%) were best explained by the separate model, and those which were, showed a high degree of overlap of the separate kernels or low rates of firing, we excluded this category from further analysis and chose the model with the next lowest BIC score for these neurons.

We defined the time modulation index as the minimum of the two estimated normalized shape parameters for each cell, with a low value near zero indicating a strong temporal modulation, and a value of one being no temporal modulation. We calculated the degree of odor modulation by computing the lifetime sparseness (Vinje and Gallant 2000).

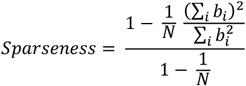

Where *b*_*i*_ is the estimated odor response vector and *N* is the number of odors (*N* = 4).

### Decoding model

For the windowed decoder, the three task factors we decoded where, the *odor* presented to the animal, the location of the *barrier* in the current block, and the upcoming *turn* decision of the animal. We used a linear discriminant decoder for these three factors. For each cell and trial, we took the average ΔF/F value in 27ms wide, non-overlapping temporal windows aligned to the start of the fixation. Odor and turn decoders were cross-validated using a n-fold random selection, where n is the number of blocks within a session. Barrier location was cross-validated using temporal partition scheme, where for each fold, a single block was selected as a test and the remaining blocks as training. This partition scheme was specifically designed to ensure the decoder was capturing barrier location signals that generalized across experimental blocks rather than block-specific characteristics of the data (Harris 2021).

To compare the performance of classifiers to chance, we constructed 400 null shuffles for each factor. For the odor and the turn factors, the trial identity was randomly permuted within the current block of trials, this ensured that the nulls reflected the local statistics of the real data. For the barrier location factor, we created a shuffled sequence of barrier location blocks that respected the original sequence (i.e. same number of each condition, and no repeats of the same barrier location on adjacent blocks). For the turn decoder we only considered trials from the two *changed turn goals*, we excluded the *fixed turn* goals because the animal’s behavior was almost entirely the correct turn for each odor, and so any odor coding could be mistaken for turn coding.

For the block lagged odor decoder, we trained a separate linear decoder for each block for the odor identity and evaluated each decoder on the test data from all other blocks; we averaged the decoder performance as a function of the block lags between training and test.

For the block identity decoder, we considered the average active of active cells in the whole trial time window, and used a multiclass categorical linear discriminant decoder. The decoder was cross validated with a 5-fold partition, where every trial in each block was randomly assigned to one of 5 equal sized partitions. This ensured the that every training and test partition had some trials from every block.

### Representational similarity analysis

For each cell and every trial within a session we calculated a population vector of the average ΔF/F response for each neuron for the whole trial following the presentation of the odor. A representational similarity/distance matrix was then constructed by taking the population vector (normalized to zero to one for each cell) Pearson’s correlation between every pair of trials. The average neural distance as a function of trial lag was calculated in bins of 20 trials. The comparison across animals for the fixed vs changed action odor/goals was made based on the first trial lag bin. Since the values in the representational similarity matrix are not independent observations, we only considered the similarity of adjacent trials for statistical testing. We took the population vector correlation between two adjacent trials and modeled it as a multiple, mixed effects regression model across the whole dataset. For the fixed vs change analysis, we selected only adjacent trials of the same odor; the fixed effects were the category of odor/goal (fixed or changed) and the absolute time difference between trials; the random effects structure included random intercepts for animal identity and session. For the turn error analysis, we selected only adjacent trials of the same odor for changed odor/goals (since animals made few errors for fixed odor/goals); the fixed effects we examined were: if the animal made a turn error on the first trial and the absolute time difference between the trials. The random effects structure included random intercepts for animal identity and session.

For the barrier location decoding analysis, we used the neural representational similarity matrix and the category similarity matrix for the closed barrier location for each trial (zero if same, one if different) We then shuffled the closed barrier location in the same manner as the linear decoding analysis, shuffling the closed barrier location ID for each block, but respecting the structure of the task and calculated a series of null category similarity matrices. Each of these was correlated to the neural representational similarity matrix to generate a distribution of correlation scores under the null that the true value could be compared to. In this way we could test for representations of closed barrier that might drift, but showed some above average similarity when comparing same and different closed barrier location blocks.

### Neural network modeling

We modelled the learning of the correct action for a given stimuli following a block change with a single layer feedforward network. The input layer represents a state input, reflective of the state representation in CA1. The input layer x projects to two output units y through a weight matrix w. The activity in the output units are the weighted sum of the activity of the input layer.

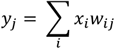

The values of the action units are passed through a softmax function, that generates a probability of taking a left or right action.

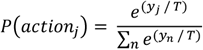

The weights were adjusted with a supervised Hebbian rule

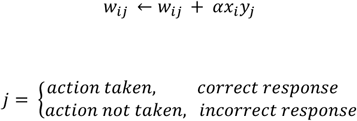

*w*_*ij*_ is the synaptic weight between the i^th^ input unit and the the j^th^ output unit, α is the learning rate. This rule corresponds to a strengthening of the synapse for the correct response regardless of the outcome. Such a rule could be implemented by eligibility traces established during action selection. After the action has been taken and the outcome evaluated, If the action taken was correct, the action taken index is potentiated, if the action was incorrect then the action not taken index is potentiated (counterfactual learning). The weights of synapses were restricted to grow no larger than 1 (the maximum value of the input units), to ensure that weights did not continue to increase indefinitely. There is no decay term nor is there any re-normalization. The was no reduction of any weights for the incorrect action whether it was taken or not.

The activity of the input layer was determined by a generative model that was based on the dynamics of the real neural data. For each simulated cell we selected a session response profile by resampling the from the temporal kernel shapes from the encoding model for real neurons and randomly generated a center point for the response within that session. This gave a predetermined response kernel which spanned the length of the simulated session and was indexed by the number of total trials in a session.

In the **time-driven** condition, the state index n of the response kernels advances by 1 on every trial:

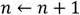

In the **error-driven** condition is only advanced following an incorrect action

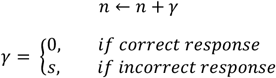

Since the total number of errors is not known in advance, the step size s must be adjusted dynamically to ensure that the full response profile is covered by the end of the session. We used the following procedure to determine s; s is initialized with a value taking into account the number of errors in real animal data to cover the full response profiles, we first run an estimate the expected number of errors by running. If the estimated step size leads to premature completion or incomplete coverage, it is iteratively refined. This ensures that the average, long run response profiles of simulated neurons in the error-driven condition match the **time-driven** condition and are representative of the neural data in both cases.

We wanted to isolate the contributions of representational drift to the learning profiles, so we restricted the simulation to cells with single odor representations (which are present in many actual cells recorded); cells were simulated with a binary one-hot response to either odor, which was then multiplied with the temporal kernel previously selected. This ensured that behavioral effects we saw in the simulation would be the result of the modulation of the drift rather than any correlation of the response vectors for each odor.

We ran the simulation with a simplified version of the task that only featured two learned odors (since the correct action for fixed odors would simply be learned initially and be maintained throughout the session), with the three closed door positions. For each session we presented 360 trials in blocks of 72 trials, this was to ensure that the network more completely leaned the rule for the current block, before having to switch to a new block. We simulated 200 sessions to generate the learning curves.

The softmax inverse temperature (*T* = 5) and learning rate (α = 0.4) were adjusted so there was a suitable match to the actual animal behavioral data. For the simulated engram reactivation condition, we re-instantiated the activity on each trial of a session through the final weight matrix *w* that was learned after training and recorded if the network generated the appropriate action for that given trial. The final engram reactivation score was the proportion of these trials that were correct.

For the Hebbian plus depression condition, we allowed the weights to adjust downwards with the same learning rate for the incorrect action. The softmax inverse temperature (*T* = 10) and learning rate (α = 0.1) were adjusted for this condition to give a similar learning profile following block changes.

