## Supplemental Material for "Error-driven changes in hippocampal representations accompany flexible re-learning"

### Supplementary figures

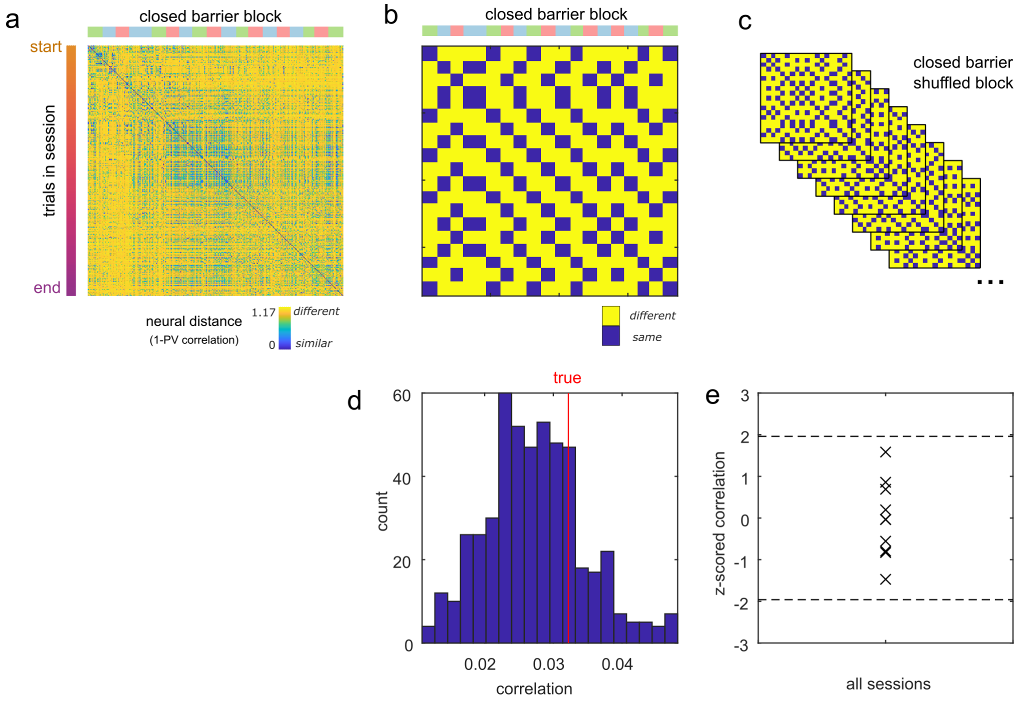


**Fig S1 – Representational similarity decoder does not detect any closed barrier block representation.** **a –** Representational similarity matrix for a single session, showing for every pair of trials, how similar the population vector of average activity was. Along the top edge is the closed barrier location block indicated with a color for each of the three barrier locations. **b –** The similarity matrix for the barrier location blocks, showing the theoretical pattern of similarity for a stable, static contextual code. **c** – Shuffling the sequence of the barrier location identity (while respecting the generative sequence) provides a set of similarity matrices under the null hypothesis of no contextual coding. **d** – the distribution of the correlation of neural representational similarity matrix with the null similarity matrices (blue) compared with the true similarity matrix (red). **e –** the correlation of neural with barrier location similarity matrices Z-scored to their own respective null distributions.

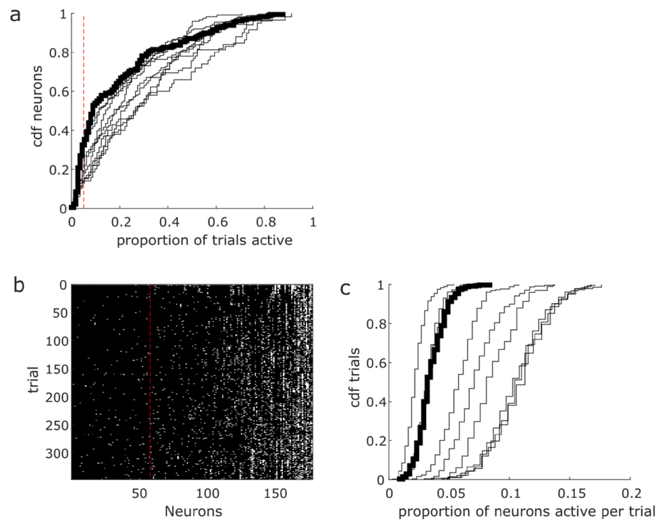


**Fig S2 – Sparse activity throughout the recording session**

**a –** The empirical cumulative distribution of the proportion of trials each neuron was active for, the red dotted line indicated the cut off of 5% of trials. Each line is an individual session, the thick line is the distribution for the session shown in b. **b –** The activity matrix for a single session, black indicates no detected transient activity for a neuron on a given trial, white indicates there was at least one transient detected. The neurons are sorted according to the proportion of trials each neuron was active for. The red dotted line indicates the threshold for inclusion in analysis. **c –** The empirical cumulative distributions for the number of neurons active on each trial. Each line is an individual session, the thick line is the distribution for the session shown in b.

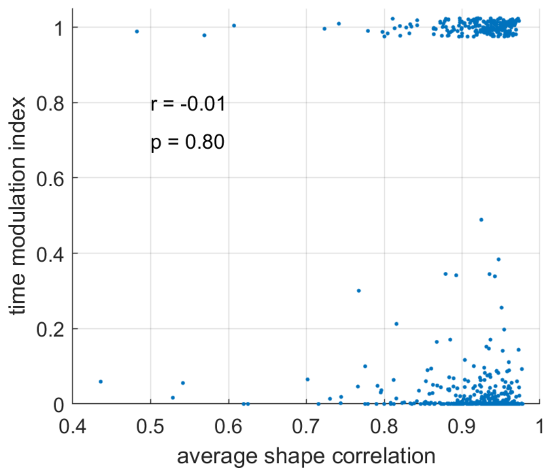


**Fig S3 – Recording stability**

Scatter plot of all cells showing the average shape correlation (a measure of the quality of the isolation quality) and the time modulation index, which is a measure of how modulated each cell was as measured with the encoding model. The numbers in the plot show the Pearson's correlation r value, and associated p-value. Vertical jitter has been added to those cells that were not temporally modulated (temporal modulation index of zero).

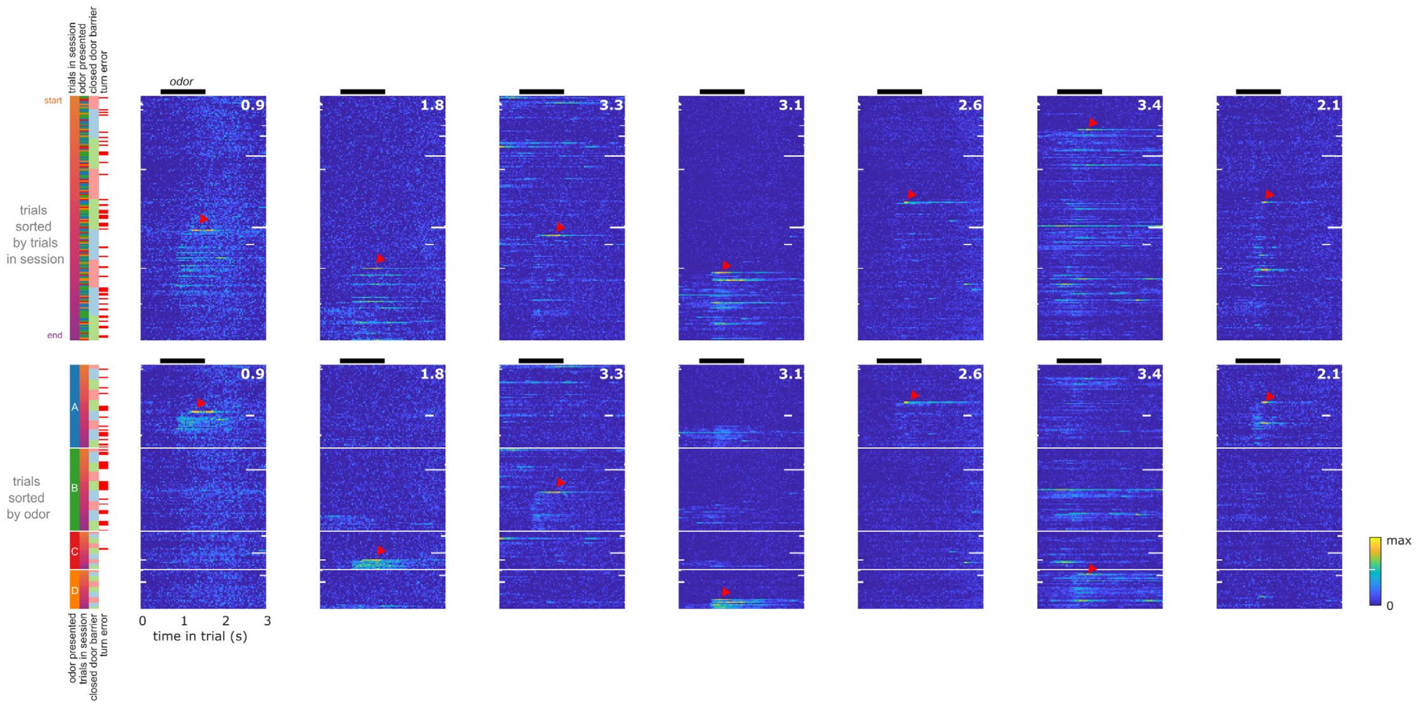


**Fig S4 – Examples of large calcium transients initiation the formation of odor response**
Here showing 7 cells in from a single session. Top row is trials sorted by the time in a session; the bottom row is the same cells sorted first by the odor delivery, and then by the time within the session. The color bar at the side shows the variables for each trial. For these neurons, a strong transient (red arrow) was seen just as the neuron began to show responsiveness to a certain odor. The strong transient also occurs later within the trial compared to the eventual odor responsive field. The inset number is the maximum dF/F of the color scale for each cell.

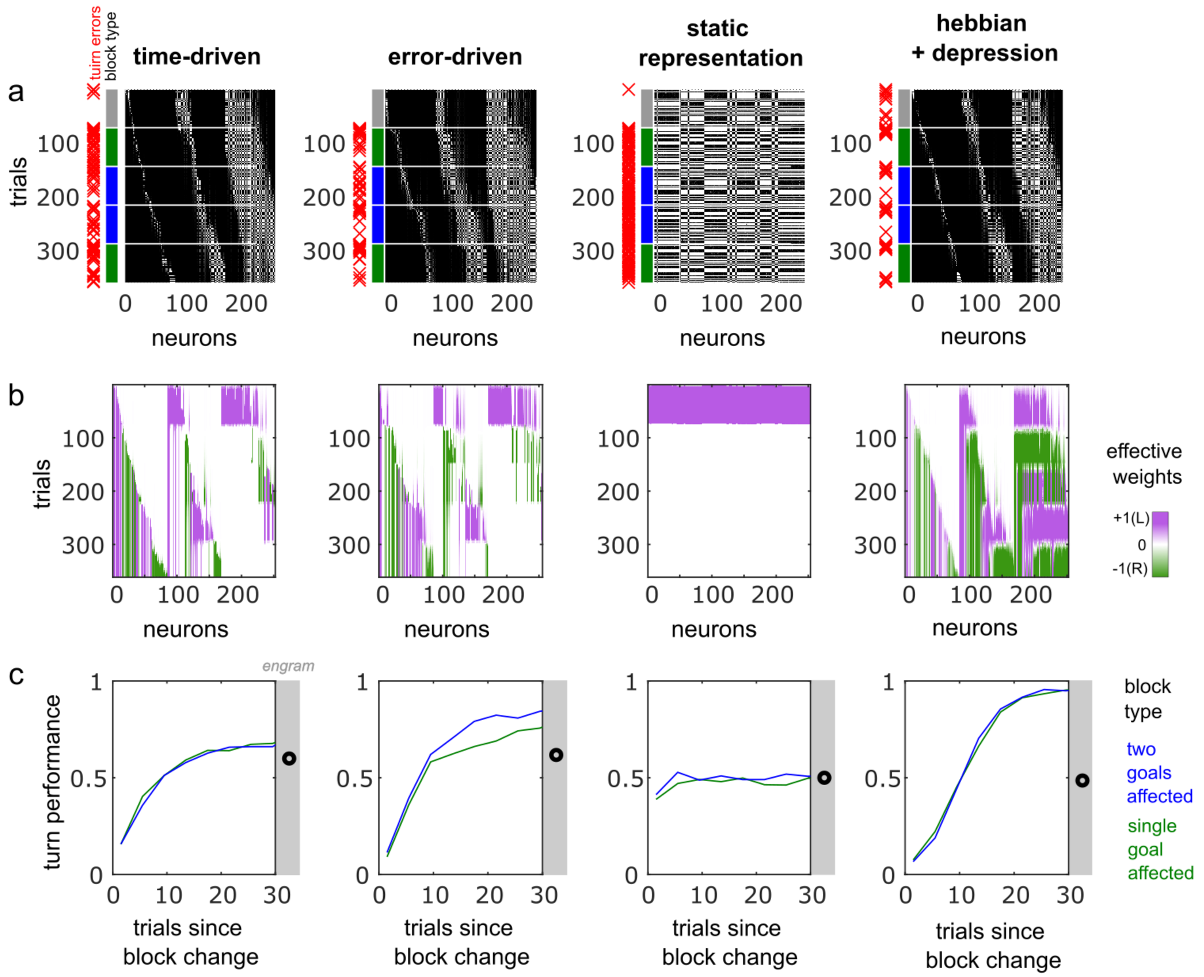


**Fig S5– Effective weights**

**a** – Simulated neural activations for a single session for four conditions: time-driven drift; error-driven drift; neural activations for the two odors were static for all trials; and the forth condition where the representation is time-driven, but the learning rule allows for depression of weights for the incorrect action. Display is the same as for Fig. 6, with neurons sorted by the total activity and then by the center of mass **b** – Effective weights for each neuron, each input neuron projects to two output neurons (L and R, Fig. 6). By combining these two weights, we can see how each neuron contributes to the decision to turn as a function of trials in the session. The effective weight for a neuron may be zero if both of the projection weights to the output neurons are the same value, effectively cancelling them out in the final decision. **c** – Performance curves for the correct turn relative to the block transition. With the given architecture, that does not allow any reduction in weights, the network is unable to adapt to the changing task demands with a static representation. When weights are allowed to be depressed, the network is able to learn effectively through the session, but does not show any above chance recall for the engram reactivation test.
